# Using protein interaction networks to identify cancer dependencies from tumor genome data

**DOI:** 10.1101/2020.08.27.270520

**Authors:** Heiko Horn, Christian Fagre, Anika Gupta, Kalliopi Tsafou, Nadine Fornelos, James T Neal, Kasper Lage

**Affiliations:** Stanley Center for Psychiatric Research, Broad Institute of MIT and Harvard, Cambridge, MA, USA; Cancer Program, Broad Institute of MIT and Harvard, Cambridge, MA, USA; Broad Institute of MIT and Harvard, Cambridge, MA, USA; Department of Biomedical Informatics, Harvard Medical School, Cambridge, MA, USA; Department of Surgery, Massachusetts General Hospital, Boston, MA, USA

## Abstract

Genes required for tumor proliferation and survival (dependencies) are challenging to predict from cancer genome data, but are of high therapeutic value. We developed an algorithm (network purifying selection [NPS]) that aggregates weak signals of purifying selection across a gene’s first order protein-protein interaction network. We applied NPS to 4,742 tumor genomes to show that a gene’s NPS score is predictive of whether it is a dependency and validated 58 NPS-predicted dependencies in six cancer cell lines. Importantly, we demonstrate that leveraging NPS predictions to execute targeted CRISPR screens is a powerful, highly cost-efficient approach for identifying and validating dependencies quickly, because it eliminates the substantial experimental overhead required for whole-genome screening.

---

Cells become tumorigenic when genomic driver events confer a positive selective advantage on them. Examples of driver events are somatic mutations, copy number alterations, or genomic fusions that promote cell growth and oncogenesis. Identifying the genes affected by driver mutations in tumors is crucial to gaining mechanistic and therapeutic insights into cancer etiology. Over the last decade, tens of thousands of tumors have been sequenced and methods such as MutSig, Oncodrive, GISTIC, RAE and MutPanning have identified hundreds of genes prone to driver events that promote tumor formation (Dietlein et al. 2020; Lawrence et al. 2014; Mermel et al. 2011; Mularoni et al. 2016). Of equal, if not greater, therapeutic interest are genes under purifying selection that when mutated negatively impact tumor development or maintenance. Some genes (e.g., PIK3CA, ESR1 and AFF1) (Ghandi et al. 2019; Ilic et al. 2017) can be both oncogenes and dependencies and are particularly attractive candidates for pharmacological intervention.

Cancer mutation rates can be modeled as randomly distributed across the tumor genome and are influenced by DNA sequence composition and genomic position, replication timing, transcription-coupled DNA damage repair and mutational hotspots (Dietlein et al. 2020; Lawrence et al. 2014; Weghorn & Sunyaev 2017). Therefore, genes under purifying selection should, in principle, be detectable from cancer genome data as those with fewer mutations than expected. However, because cancer genome data are both noisy (e.g., due to mutation rate heterogeneity, germline variants in sequencing data, gene copy number and mutation rates) and sparse (most genes are not mutated), measuring a statistical signal for genes under purifying selection requires large sample sizes (Tsherniak et al. 2017; Weghorn & Sunyaev 2017).

Despite these challenges, genes under purifying selection have been successfully identified by constraining their search to highly specific features of the cancer genome. These include genes that introduce a vulnerability through the loss of one allele as a result of genomic copy number changes (e.g., CYCLOPS) (Nijhawan et al. 2012) and functionally coherent gene sets derived from The Cancer Genome Atlas with mutually exclusive loss-of-function signatures (Ryan et al. 2014). While important and pioneering, none of these approaches can be applied in an unconstrained and genome-wide manner to directly detect cancer vulnerabilities from the tens of thousands of available cancer genomes.

Consequently, cancer researchers have historically executed small-scale gene perturbation experiments in cancer cell lines to assess the resulting effects on cell growth (Bass et al. 2009; Etemadmoghadam et al. 2010; Garraway & Lander 2013; Kim & Sabatini 2004; Lopez & Hanahan 2002; Mansouri et al. 1998; Okhrimenko et al. 2005; Ramsay & Gonda 2008). More recently, the development of genome-wide RNAi knockdown and CRISPR-Cas9 knockout experiments has made it possible to systematically study all genes in a wide range of cell lines (Cheung et al. 2011; Meyers et al. 2017; Tsherniak et al. 2017; Wang et al. 2015). These screens have identified many pan-essential and cell line-specific dependencies that can be reproducibly found in independent screens (Dempster et al. 2019; Meyers et al. 2017) and it is now possible to personalize these methods to identify unique, patient-specific dependencies on a treatment-relevant timescale (Hong et al. 2016). Despite this tremendous progress, the identification of all possible dependencies would require screening more than 5,000 cell lines, an extremely resource- and time-consuming endeavor (Tsherniak et al. 2017). Therefore, a computational method that can reliably identify genes under purifying selection directly from cancer genome data is highly relevant to oncology.

We previously developed NetSig, a robust statistic that combines cancer mutation data with protein network information to discover cancer driver genes (Horn et al., 2018). Here, we used an analogous approach to create Network Purifying Selection (NPS), a statistic that aggregates weak signals of negative selection across a gene’s first order, high-confidence, protein-protein interaction (PPI) network and we show that this signature is predictive of the gene itself being under purifying selection. To calculate the signal of purifying selection for a given gene, we considered the first order interactors of its encoded protein (hereafter designated as neighborhood) from PPI evidence (InWeb3 and InBio Map (Lage et al. 2007; Li et al. 2017), see Online Methods) integrated with cancer genome data (Lawrence et al. 2014) (**Fig. 1a**). We calculated a gene-specific NPS statistic using the genetic data from 4,742 human cancers across 21 cancer types and obtained a set of 91 gene dependencies that we call NPS5000 (as it consists of about 5,000 patient samples and to follow the naming convention of previous work (Horn et al. 2018; Lawrence et al. 2014)) (**Supplementary Table 1**).

**Figure 1.**
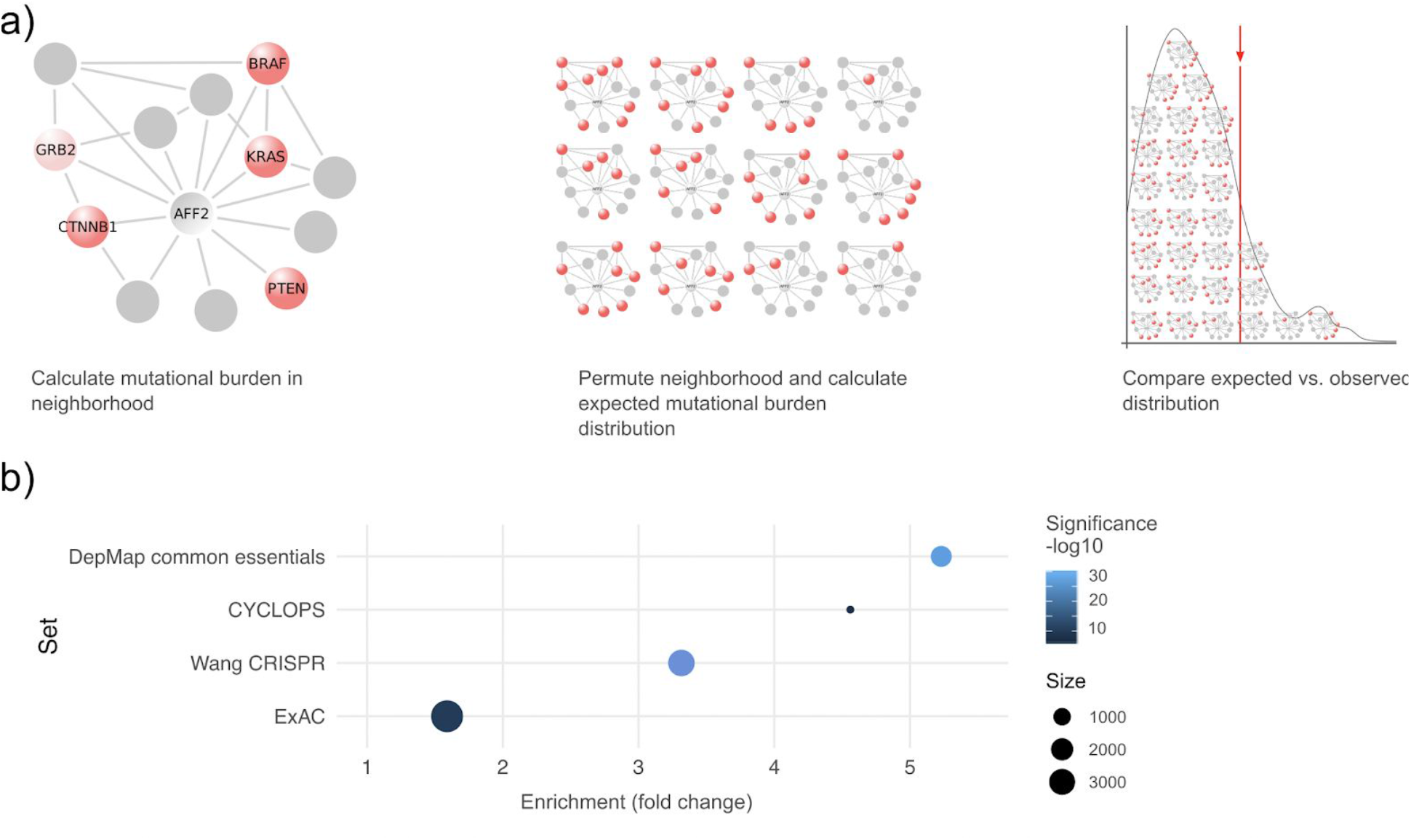
Network Purifying Selection (NPS) applied to cancer dependency data sets. **a)** Study overview depicting the calculation of the mutation burden in the neighborhood of a candidate cancer dependency gene product, AFF2, and use of a permutation scheme to empirically derive the significance of the aggregated mutation burden across its interaction network. **b)** Enrichment of expected vs. observed mutation burden fold change of candidate genes from four dependency data sets.

To account for knowledge contamination, we used a permutation scheme that considers the network topology of the neighborhood. Briefly, this strategy consists in replacing every neighbor of a given gene with a gene having a similar connectivity in the network (see **Methods** for more details) and calculate the mutation burden of this permuted neighborhood. We determined the predictive performance of the NPS algorithm on four well-established gene dependency data sets (**Fig. 1b**). First, we applied NPS to CYCLOPS (Copy-number alterations Yielding Cancer Liabilities Owing to Partial losS) and found a strong enrichment of the NPS5000 gene set (fold change = 4.6; *P* < 0.01, hypergeometric test) (Nijhawan et al. 2012). Second, we examined ExAC, a human population genetic data set of ∼80,000 exomes (pLI > 0.9) and found a significant NPS5000 enrichment (fold change = 1.6, *P* < 0.001, hypergeometric test) (Lek et al. 2016). The third data set used for assessing the NPS predictions was obtained from a small-scale gene knockout experiment in cancer cell lines using CRISPR-Cas9 and Gene-trap technologies (hereafter referred to as the Wang-CRISPR data set (Wang et al. 2015)). We found strong enrichment of our candidate genes (fold change = 3.3, *P* < 4.3e-23, hypergeometric test) as expected from a study focusing on genes that are essential to cell function. Lastly, we looked at dependency probabilities in one of the largest gene knockout studies available, the Cancer Dependency Map project (DepMap 2020), a genome-wide CRISPR-Cas9 knockout in 769 cell lines, and here too found strong enrichment of the NPS5000 dependency gene set (fold change = 5.23, *P* = 3.73-28, hypergeometric test) (Tsherniak et al. 2017).

To determine whether the NPS algorithm is predicting pan-essential genes or nominating potential cancer cell line-specific dependencies, we first stratified the NPS5000 candidate genes by their ExAC pLI score (pLI > 0.9), which reflects the tolerance of a given gene to the loss of function and can therefore be seen as representative of the evolutionary constraints acting on genes. We next used the DepMap Avana 20Q2 data set to confirm that the 91 NPS-identified genes are pan-essential (**Supplementary Table 2**). Strikingly, genes classified as dependencies in the ExAC data set tend to also be pan-dependencies in cell lines (*P* = 2.93e-05, Wilcoxon signed-rank test), indicating that NPS is able to predict pan-essential genes from tumor sequencing data. Based on both the ExAC pLI and calculated dependency scores, we selected 58 genesand 40 predicted pan-dependent gene controls for experimental validation. We tested the 58 candidate genes for essentiality in tumor development in a small scale experiment using the DepMap CRISPR-Cas9 knockout protocol. We selected six cell lines that are representative of diverse tissues (see **Methods** for more details) and of which three have since been tested in the DepMap project, meaning that our validation data has been replicated in another study and contains unique cell types. We analyzed the screening results using CERES to account for sgRNA multi-target effects (Meyers et al. 2017)(**Supplementary Tables 3 & 4**). Compared with DepMap results, our candidate genes displayed similar behavior in the three tested cell lines (A375, G401, PANC1) (**Fig. 2a,b**).

**Figure 2.**
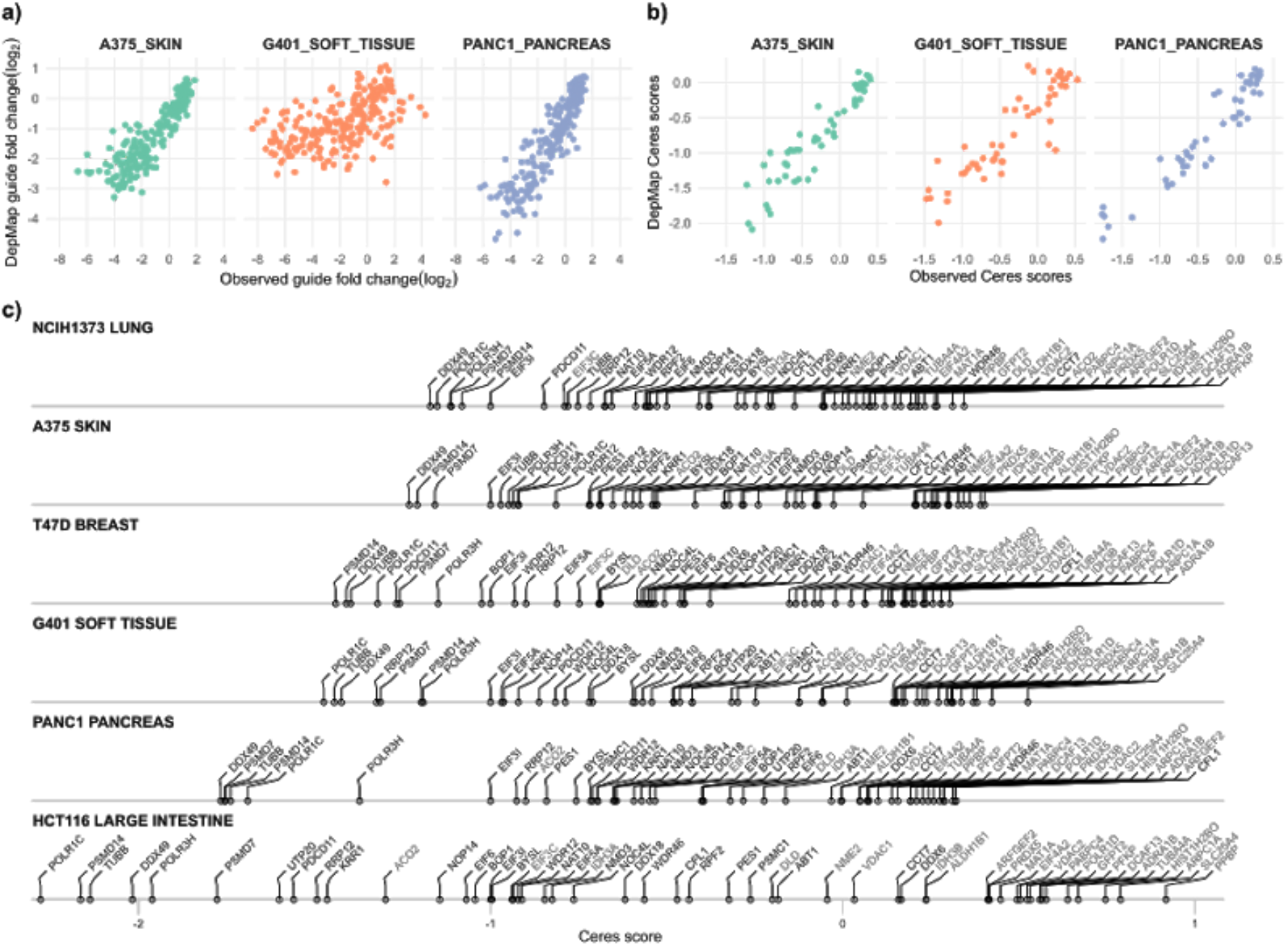
*In vitro* validation of Network Purifying Selection (NPS)-predicted cancer dependencies. **a,b)** Comparison of sgRNA effects on gene expression (fold change) (a) and CERES gene scores (b) in DepMap and CRISPR Dropout data of 58 genes in three cell lines. Dots represent each of the 58 genes. **c)** CERES gene scores (copy number corrected gene scores) for all genes in the 6 tested cell lines.

Our results show that NPS identifies cancer dependencies from tumor sequencing data alone using a network-based approach that excludes mutation information of the gene being tested, this way preventing biases introduced by the mutation rate of the index gene. Enrichment analyses comparing the NPS-imputed genes with CYCLOPS and CRISPR-Cas9 cancer cell-essential genes reveal significant NPS-enrichment, indicating that a substantial subset of the genes identified through this network approach are known cancer dependencies. The overlap between NPS-imputed genes and genes in the ExAC database indicates that the NPS method has identified genes whose integrity is essential to normal cell function, supporting that other genes significantly identified in NPS but not in ExAC are likely to be tumor-specific dependencies. In addition, genes predicted by NPS lead to statistically significant tumor growth aberrations when knocked out (as determined in DepMap, Meyers et al., 2017), showing the direct relevance of these genes in sustaining tumor proliferation. Importantly, we demonstrate that leveraging NPS predictions to execute targeted CRISPR screens is a powerful and highly cost-efficient approach for identifying and validating dependencies quickly, because it eliminates the substantial experimental overhead required for whole-genome screening.

Our approach corroborates previous studies on cancer dependencies while also identifying a novel set of potential cancer vulnerabilities. Importantly, we illustrate that it is possible to identify *bona fide* genes under purifying selection directly from cancer genome data. We expect that NPS will be widely applicable as more cancer genomes are sequenced. The NPS code is available at www.lagelab.org, and we make the latest predictions using an updated network and mutational data available as a community resource.

## Methods

### Calculating the network purifying selection score

For a given index gene, the Network Purifying Selection statistic is formalized into a probabilistic score that reflects the index-gene-specific composite purifying selection (i.e., the aggregate of single-gene MutSig suite Q values reported in (Lawrence et al. 2014)) across its first order biological network and is calculated via a three-step process. First, we identify all genes it interacts with directly at the protein level, only including high-confidence quality-controlled data from the functional human networks InWeb3 and InBio_Map (where the vast majority of connections stem from direct physical interaction experiments at the protein level). Second, the composite purifying selection score across members of the resulting network is quantified by aggregating single-gene MutSig suite Q values from Lawrence et al. into one value <l using an approach inspired by Fisher’s method for combining P-values: cp∼-2kLi=0ln(qi) where q is the MutSig suite Q value for gene i, and k is the number of genes in the first order network of the index gene (i.e., the index gene’s degree). Third, by permuting the neighborhood using a node permutation scheme, we compare the aggregated burden of mutations to a random expectation. In this step, the degree of the index gene, as well as the degrees of all genes in the index gene’s network is carefully considered to be of similar connectivity. The final NPS score of an index gene is therefore an empirical P value that reflects the probability of observing a lower composite mutation burden than expected across its first order physical interaction partners (at the protein level) normalized for the degree of the index gene as well as the degrees of all of its first order interaction partners. Because we are interested in estimating the purifying selection independent of the index gene, this gene is not included in the analysis and it does not affect the NPS calculation.

### Generating the NPS5000 set

NPS probabilities were determined for every gene in InWeb 3 and InWeb_IM that was covered by interaction data using 10^6 permutations. The FDR Q values were calculated as described by Benjamini and Hochberg (Benjamini & Hochberg 1995), based on the nominal P values. We saw a correlation between NPS significance and the number of interactors of a given gene (R=0.39, Pearson) (**Supplementary Figure 1**). We calculated the consensus candidate gene set across 4,742 human cancer genomes by taking the union of all nominally significant genes from InWeb 3 and all significant genes after correction for multiple testing from InWeb_IM, which led to a set of 91 genes namedNPS5000 and used in all downstream analyses.

### Dissecting tumor gene essentiality

We tested the effects of NPS gene knockouts in four different data sets: 1) The CYCLOPS (Copy-number alterations Yielding Cancer Liabilities Owing to Partial losS) data is induced haploinsufficiency due to copy number loss. We defined this set as genes in their manuscript (Nijhawan et al. 2012). 2) ExAC (Exome Aggregation Consortium) a source of about 80,000 exome data sets. We selected the set of genes with pLI < 0.9 (Lek et al. 2016), indicating that these genes are under evolutionary constraint. 3) The Wang-CRISPR data set of gene knockouts in cell lines using both CRISPR and Gene trap (Wang et al. 2015). Defined dependency genes as genes in their supplement. 4) The Dependency Map dataset version 20Q2 (DepMap 2020) of cancer cell line dependencies.

To select genes for downstream validation, we selected genes that had a > 50% probability of being a dependency in any cell line in the Avana data and a pLI < 0.9 in ExAC (to deplete non-cancer-specific dependencies).

### Cell Lines

PANC1 and A375 cells were cultured in Dulbecco’s modified eagle medium (DMEM, Gibco), NCIH1373 cells were cultured in RPMI 1640 medium (Gibco), HCT166 cells were cultured in McCoy’s 5A medium (Gibco), and T47D cells were cultured in phenol red-free RPMI 1640 medium (Gibco). All culture media was supplemented with 10% (v/v) fetal bovine serum (Sigma-Aldrich), 100 units/mL penicillin, 100 ug/mL streptomycin, and 2 mM L-glutamine (Gibco). Cells maintained at 37°C in 5% CO2, and passaged 2-3x per week to maintain confluence between 20% and 80%. All cell lines sourced from ATCC.

### CRISPR-Cas9 Dropout Screen

Cas9-expressing cell lines were generated by lentiviral transduction of a Cas9 expression vector with a blasticidin resistance cassette (pLX_311-Cas9, Addgene #96924). Cells were transduced by addition of viral media in the presence of 8 ug/mL polybrene (Santa Cruz Biotechnology), centrifuged at 1178 x g for 30 minutes at 37C, removing viral media after 16 hours. Transduced cells were selected by addition of 5-15 ug/mL blasticidin (Life Technologies, Inc) two days post-transduction, and cultured for at least 14 days before screening. A library of 256 CRISPR sgRNA targeting 58 genes was cloned into the CROP-Seq-Guide-Puro expression vector (Addgene #86708) and packaged into lentivirus, then transduced as described above into respective Cas9 expressing cell lines across 3 replicates at a MOI<1, and at a representation of at least 1 x 10^3 cells per guide. Two days after transduction, cells were selected for 48 hours with puromycin (Life Technologies, Inc) at 2-4 ug/mL. Cells were maintained at a density of at least 0.3 x 10^6 to maintain library representation for 21 days (passaging every 3-4 days), harvesting at least 0.3 x 10^6 cells per replicate by centrifugation at 4, 14, and 21 days post-transduction.

### Next-Generation Sequencing

Cells are washed once with PBS, pelleted, and re-suspended in lysis buffer (1 mM CaCl2, 3 mM MgCl, 1mM EDTA, 1% Triton X-100 (Sigma-Aldrich), 10 mM TrisHCl, pH 7.5) at a concentration of 10,000 cells/uL. Lysates were incubated at 65C for 10 minutes, then 95C for 15 minutes. Genomic regions containing guide sequences are PCR amplified using paired primers with 25 uL 2x JumpStart Taq Polymerase Ready Mix (Sigma-Aldrich), 1.5 uL 10 uM primer pair (CropSeq_NGS_P7, CropSeq_NGS_P5), 12.5 uL cell lysate, dH20 up to 50 uL at the following cycling conditions: 95C for 5 min, 28 cycles at 95C for 20 s, 55C for 30s, 72C for 30s, and 72C for 4 minutes. Individual samples are indexed with a second PCR step using paired 3’ and 5’ Illumina TruSeq indexing primers, using 12.5 uL JumpStart Taq Polymerase Ready Mix, 1.25 uL 5 uM index primer pair, 1.25 uL product from first PCR, and dH20 to 25 uL at the following cycling conditions: 95C for 5 mins, 18 cycles of 95C for 20s, 55C for 30s, 72C for 30s, and 72C for 4 minutes. PCR products are pooled at equal volume ratios and run on an agarose gel, gel purified, and quantified using a QuBit 4 Fluorometer (Invitrogen). Samples sequenced on a Miniseq (Illumina) using standard Illumina protocols with a 20% PhiX spike-in.

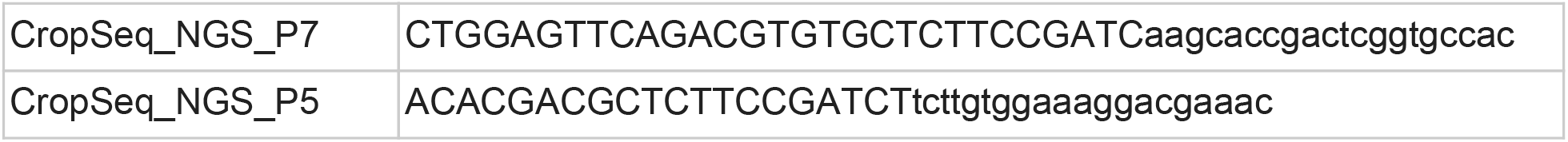

### Screen Data Analysis

Reads are deconvoluted by unique 5’ and 3’ index pairs from raw sequencing using the standard MiniSeq pipeline. For each replicate of one same sample, reads containing the expected guide sequence were counted and normalized against total reads from that sample. CERES scores were calculated as described in (Meyers et al. 2017). We did not transform CERES scores into probabilities (as is standard in recent Dependency Map publications) because our data set was too small to satisfy score calibration requirements.

## Supporting information

Supplementary Table 1 - NPS results

Supplementary Table 2 - NPS candidates

Supplementary Table 3 - Raw read counts for validation

Supplementary Table 4 - Ceres scores from validation

## Bibliography

Bass AJ, Watanabe H, Mermel CH, Yu S, Perner S, et al. (2009). SOX2 is an amplified lineage-survival oncogene in lung and esophageal squamous cell carcinomas. Nat. Genet. 41(11):1238–42

Benjamini Y, Hochberg Y. (1995). Controlling the false discovery rate: A practical and powerful approach to multiple testing. Journal of the Royal Statistical Society: Series B (Methodological). 57(1):289–300

Cheung HW, Cowley GS, Weir BA, Boehm JS, Rusin S, et al. (2011). Systematic investigation of genetic vulnerabilities across cancer cell lines reveals lineage-specific dependencies in ovarian cancer. Proc Natl Acad Sci USA. 108(30):12372–77

Dempster JM, Pacini C, Pantel S, Behan FM, Green T, et al. (2019). Agreement between two large pan-cancer CRISPR-Cas9 gene dependency data sets. Nat. Commun. 10(1):5817

DepMap B. (2020). DepMap 20Q2 Public. Figshare

Dietlein F, Weghorn D, Taylor-Weiner A, Richters A, Reardon B, et al. (2020). Identification of cancer driver genes based on nucleotide context. Nat. Genet. 52(2):208–18

Etemadmoghadam D, George J, Cowin PA, Cullinane C, Kansara M, et al. (2010). Amplicon-dependent CCNE1 expression is critical for clonogenic survival after cisplatin treatment and is correlated with 20q11 gain in ovarian cancer. PLoS ONE. 5(11):e15498

Garraway LA, Lander ES. (2013). Lessons from the cancer genome. Cell. 153(1):17–37

Ghandi M, Huang FW, Jane-Valbuena J, Kryukov GV, Lo CC, et al. 2019. Next-generation characterization of the Cancer Cell Line Encyclopedia. Nature. 569(7757):503–8

Hong AL, Tseng Y-Y, Cowley GS, Jonas O, Cheah JH, et al. (2016). Integrated genetic and pharmacologic interrogation of rare cancers. Nat. Commun. 7:11987

Horn H, Lawrence MS, Chouinard CR, Shrestha Y, Hu JX, et al. (2018). NetSig: network-based discovery from cancer genomes. Nat. Methods. 15(1):61–66

Ilic N, Birsoy K, Aguirre AJ, Kory N, Pacold ME, et al. (2017). PIK3CA mutant tumors depend on oxoglutarate dehydrogenase. Proc Natl Acad Sci USA. 114(17):E3434–43

Kim DH, Sabatini DM. (2004). Raptor and mTOR: subunits of a nutrient-sensitive complex. Curr. Top. Microbiol. Immunol. 279:259–70

Lage K, Karlberg EO, Storling ZM, Olason PI, Pedersen AG, et al. (2007). A human phenome-interactome network of protein complexes implicated in genetic disorders. Nat. Biotechnol. 25(3):309–16

Lawrence MS, Stojanov P, Mermel CH, Robinson JT, Garraway LA, et al. (2014). Discovery and saturation analysis of cancer genes across 21 tumour types. Nature. 505(7484):495–501

Lek M, Karczewski KJ, Minikel EV, Samocha KE, Banks E, et al. (2016). Analysis of protein-coding genetic variation in 60,706 humans. Nature. 536(7616):285–91

Li T, Wernersson R, Hansen RB, Horn H, Mercer J, et al. (2017). A scored human protein-protein interaction network to catalyze genomic interpretation. Nat. Methods. 14(1):61–64

Lopez T, Hanahan D. (2002). Elevated levels of IGF-1 receptor convey invasive and metastatic capability in a mouse model of pancreatic islet tumorigenesis. Cancer Cell. 1(4):339–53

Mansouri A, Chowdhury K, Gruss P. (1998). Follicular cells of the thyroid gland require Pax8 gene function. Nat. Genet. 19(1):87–90

Mermel CH, Schumacher SE, Hill B, Meyerson ML, Beroukhim R, Getz G. (2011). GISTIC2.0 facilitates sensitive and confident localization of the targets of focal somatic copy-number alteration in human cancers. Genome Biol. 12(4):R41

Meyers RM, Bryan JG, McFarland JM, Weir BA, Sizemore AE, et al. (2017). Computational correction of copy number effect improves specificity of CRISPR-Cas9 essentiality screens in cancer cells. Nat. Genet. 49(12):1779–84

Mularoni L, Sabarinathan R, Deu-Pons J, Gonzalez-Perez A, Lopez-Bigas N. (2016). OncodriveFML: a general framework to identify coding and non-coding regions with cancer driver mutations. Genome Biol. 17(1):128

Nijhawan D, Zack TI, Ren Y, Strickland MR, Lamothe R, et al. (2012). Cancer vulnerabilities unveiled by genomic loss. Cell. 150(4):842–54

Okhrimenko H, Lu W, Xiang C, Hamburger N, Kazimirsky G, Brodie C. (2005). Protein kinase C-epsilon regulates the apoptosis and survival of glioma cells. Cancer Res. 65(16):7301–9

Ramsay RG, Gonda TJ. (2008). MYB function in normal and cancer cells. Nat. Rev. Cancer. 8(7):523–34

Ryan CJ, Lord CJ, Ashworth A. (2014). DAISY: picking synthetic lethals from cancer genomes. Cancer Cell. 26(3):306–8

Tsherniak A, Vazquez F, Montgomery PG, Weir BA, Kryukov G, et al. (2017). Defining a cancer dependency map. Cell. 170(3):564-576.e16

Wang T, Birsoy K, Hughes NW, Krupczak KM, Post Y, et al. (2015). Identification and characterization of essential genes in the human genome. Science. 350(6264):1096–1101

Weghorn D, Sunyaev S. (2017). Bayesian inference of negative and positive selection in human cancers. Nat. Genet. 49(12):1785–88

