## Supplementary Figure 1 for "Using protein interaction networks to identify cancer dependencies from tumor genome data"

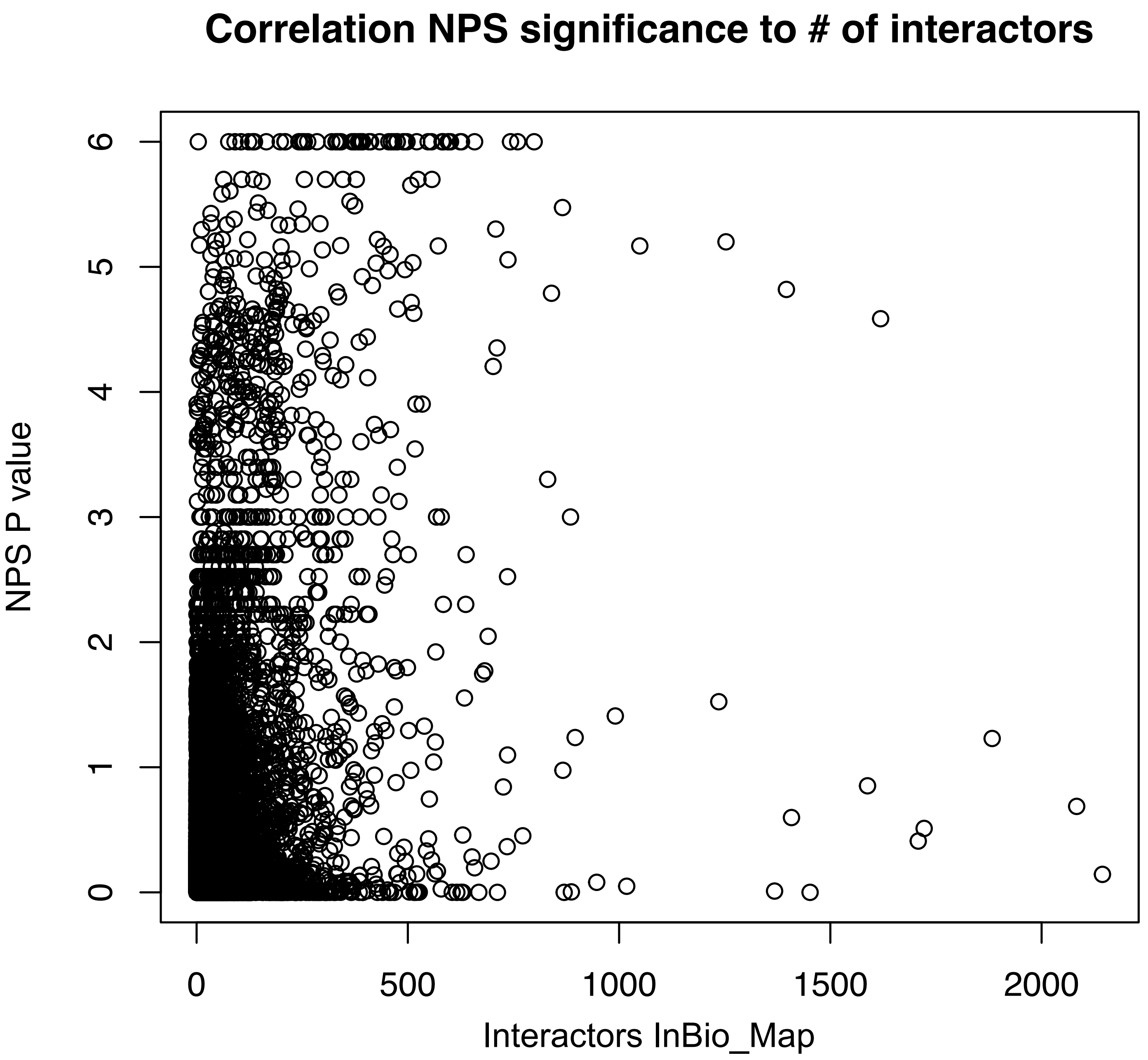


**Supplementary Figure 1**. Correlation between the number of interactors of a gene in InBio_Map and its NPS P value. The NPS P value is limited to 6 as we did a maximum of 1,000,000 iterations in the permutation step.
